# A formula for calculating 30-item Geriatric Depression Scale (GDS-30) scores from the 15-item version (GDS-15)

**DOI:** 10.1101/2022.09.22.509012

**Authors:** Yuge Zhang, Marco Hoozemans, Mirjam Pijnappels, Sjoerd M. Bruijn

## Abstract

The Geriatric Depression Scale with 30 items (GDS-30) and with 15 items (GDS-15) are both valid tools for assessing depression in older adults, but their absolute values are not directly comparable. Here, we used a dataset (n=431) with GDS-30 scores from a project concerning fall-risk assessment in older adults (FARAO) to develop and validate a formula which can be used to convert GDS-15 scores into GDS-30 scores. We found that the GDS-15 score cannot simply be multiplied by 2 to obtain the GDS-30 scores and that estimations of GDS-30 from GDS-15 are not affected by age, sex and MMSE. Therefore, the optimal formula to estimate the GDS-30 score from the GDS-15 score was: GDS-30_estimated = 1.57 + 1.95 × GDS-15. This formula yielded an estimate of GDS-30 with an explained variance of 79%, compared to 63% when GDS-15 was simply multiplied by 2. Researchers that have used the GDS-15 and want to compare their outcomes to other studies that reported only the GDS-30 are advised to use this formula.

## 1. Introduction

Our society is ageing, and it is well known that ageing comes with physical and mental problems. For instance, the prevalence of depression in older adults is around 28.4% worldwide (Hu and others 2022), with ranges of 10% to 30% in hospitals and nursing homes (Mitchell and others 2010b). The Geriatric Depression Scale with 30 items (GDS-30), designed by Yesavage and others (1983), has been shown to be a valid and reliable tool for assessing both major and minor depression in nursing home patients (Jongenelis and others 2005), primary care (Mitchell and others 2010a) and medical settings (Mitchell and others (2010b). The GDS-30 is a questionnaire, consisting of 30 yes/no items, which asks about depressive feelings over the past week. For most of these items, a “yes” answer indicates higher depressive symptoms. A sum-score of 10 is the cut-off between normal and depression and a sum-score of 20 is the dividing line between mild and severe depression. In 1986, a shortened version of the GDS-30 was proposed by Sheikh and Yesavage (1986) with 15 selected items (from the GDS-30) that had the highest correlation with depressive symptoms. The GDS-15 was also found to have good sensitivity and specificity, among older adult psychiatric inpatients (Mitchell and others 2010a; Mitchell and others 2010b), so it is a good substitute for the GDS-30 (Lesher and Berryhill 1994).

In a recent study, we came across a situation where we had administered the GDS-15, whereas the literature we were comparing our results to mostly reported results in terms of the GDS-30. The simplest solution to being able to compare our GDS-15 results to those who used the GDS-30 would have been multiplying the GDS-15 sum scores by 2. However, it was unclear if these scales are directly comparable as, to the best of our knowledge, there is no literature which provides a valid formula to ‘convert’ GDS-15 scores to equivalent GDS-30 scores that goes beyond the ‘multiply by 2’ method. Considering all items in GDS-15 come from the more extensive GDS-30, GDS-15 scores can be obtained from completely administered GDS-30 scores. Thus, here, we used a previously collected GDS-30 dataset from a community living sample to compare the validity of multiplying the GDS-15 sum scores by 2 to get GDS-30 sum scores with the validity of a linear conversion formula from GDS-15 to GDS-30 scores.

## 2. Methods

### 2.1 Data collection

The dataset we used was from a project concerning fall-risk assessment in older adults (FARAO) performed at the Vrije Universiteit, Amsterdam (van Schooten and others 2015). Participants were independently living individuals living in Amsterdam and surroundings and were included if their ages were between 65 and 99 years, and their mini mental state exam score (MMSE) was between 19 and 30. All items of the GDS-15 were contained in GDS-30, which participants filled in. The medical ethical committee of the Vrije Universiteit medical center approved the protocol (#2010/290) and all participants provided informed consent. From the individual scores on the GDS-30 items, the GDS-30 and GDS-15 scores were calculated by summing the scores of the items (after reversing the scores of the items for which this was required) according to Yesavage and others (1983) and Kurlowicz and Greenberg (2007), respectively.

### 2.2 Statistical analysis

Statistical analysis was performed in MATLAB R2021a (MathWorks, U.S.). We used linear regression to determine the relationship between GDS-30 sum score (outcome variable) and GDS-15 sum score (predictor variable). To test whether the obtained model was better than simply multiplying the GDS-15 scores by 2, we checked the goodness of fit, which was evaluated by the coefficient of determination (R^2^) and measured the error of the model in predicting the GDS-30 by root mean square error (RMSE, with lower RMSE values indicating smaller errors)(Hyndman and Koehler 2006). Moreover, we evaluated whether the intercept of the obtained model was significantly different from 0 (i.e., 95% confidence interval (CI) did not include zero), and whether the regression coefficient of the GDS-15 score as predictor variable was significantly different from 2 (i.e., the 95% confidence interval did not include 2). Age, sex and MMSE score were explored for interaction, as it might be expected that these variables affect the relationship between GDS-15 scores and GDS-30 scores, i.e. that this relationship is different for different levels of the potential effect modifiers.

## 3. Results

### 3.1 Demographic characteristics

From the FARAO cohort, 431 participants satisfied the inclusion criteria and filled in all items of GDS-30 questionnaires (see table 1). The mean age was 76.4 (SD 7.5) years, the mean GDS-30 sum score was 5.4 (4.7), and the mean GDS-15 sum score was 2.0 (2.1). This latter result already suggests that GDS-15 scores cannot simply be multiplied by 2 to get GDS-30 scores.

**Table 1.** Demographic Characteristics.

| Characteristics | Mean (SD) |
| --- | --- |
| Age (years) | 76.4(7.5) |
| Sex (% male) | 41.5% |
| MMSE | 27.4(2.4) |
| GDS-30 scores | 5.4 (4.7) |
| GDS-15 scores | 2.0 (2.1) |

### 3.2 Prediction of GDS-30 scores from GDS-15 scores

Age, sex and MMSE score showed no significant interaction with GDS-15 scores in the prediction of GDS-30 scores (see supplementary table 1). Therefore, only GDS-15 was considered as the single predictor variable of GDS-30 in the regression model. The linear regression model to estimate GDS-30 from GDS-15 was GDS-30_estimated = 1.57 + 1.95 × GDS-15. The R^2^ of the linear model predicting GDS-30 from GDS-15 was 0.79, and RMSE was 2.13 (see figure 1; marker size represents the number of subjects for that data point). When simply multiplying GDS-15 scores by 2, the R^2^ was 0.63, and RMSE was 2.59, indicating that our linear model was better than simply multiplying GDS-15 scores by 2. The intercept was significantly different from zero, (95% CI [1.29,1.84]), but the regression coefficient of GDS-15 was not significantly different from 2 (95% CI [1.86,2.05]).

**Figure 1.**
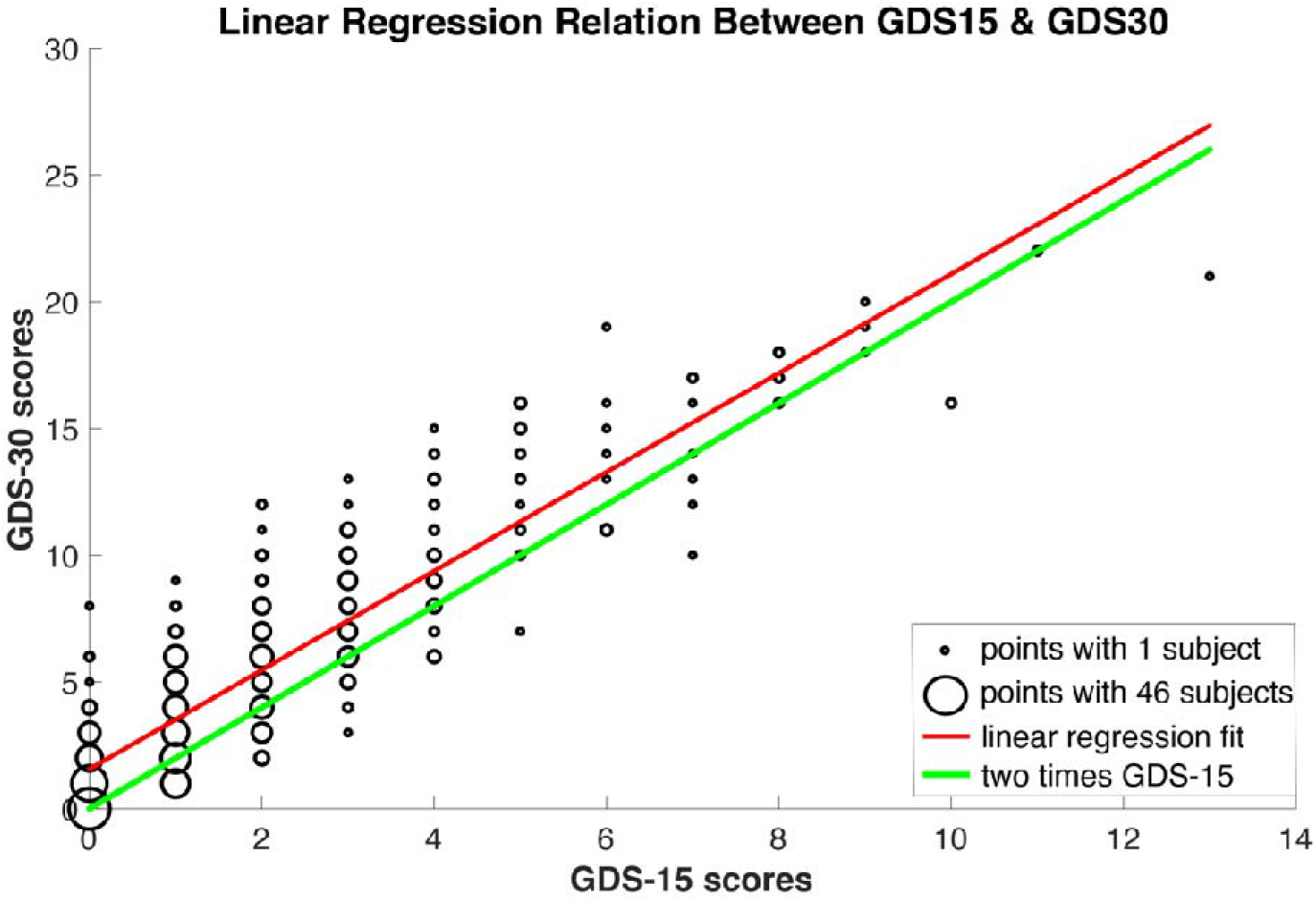
Linear regression relation between GDS-15 & GDS-30 score; marker size represents the number of subjects for that data point

## 4. Discussion

We set out to find a formula to estimate equivalent GDS-30 scores from GDS-15. In our cohort of over 400 older adults, we found the formula GDS-30=1.57 + 1.95 × GDS-15, which yielded better performance than simply multiplying GDS-15 scores by 2, as shown by the higher explained variance (R^2^) and lower error (RMSE). Although the slope was not significantly different from 2, the intercept was statistically higher than zero. This indicates that GDS-30 scores calculated using the GDS-15 times 2 method systematically underestimate the better fitting linear model by 1.6 points, which is also clearly visible in figure 1. Our analyses indicated that GDS-30 scores calculated from GDS-15 scores by using the suggested formula have a better goodness of fit and lower error than simply multiplying a GDS-15 score by two. In addition, the formula appeared to be robust as the relationship was not affected by age, sex and MMSE score. The formula can be useful when comparing the geriatric depression scale among different studies.

Our study has some limitations. First, as the data were collected in a community-dwelling sample, the validity of our formula in a more severely depressed sample remains to be tested. Second, although using non-integer estimates of the GDS-30 to compare with other studies is not a problem, for clinical interpretation, integer values are probably preferred. Lastly, our linear regression leads to a subject with score 0 on the GDS-15 having a score of 1.57 for the estimated GDS-30. While this may seem odd, it should be kept in mind that this is still well below the cut-off for depression (which is 10).

## 5. Conclusion

In conclusion, the linear regression formula developed in our study can be used effectively for estimating GDS-30 scores from GDS-15 assessments.

## Acknowledgements

Many thanks to Kimberley van Schooten of Neuroscience Research Australia, University of New South Wales, and Roel Weijer of the Department of Neurology, Leiden University Medical Center, who collected the data. YZ was funded by the China Scholarship Council (CSC) (202009110145). MP and SMB were funded by a grant from the Dutch Organization for Scientific Research (NWO) (no. 91714344 and 016.Vidi.178.014, respectively).

## Author contribution

YZ contributed to formal analysis, software, funding acquisition, writing the original draft, reviewing and editing of the draft. MH contributed to formal analysis, methodology, reviewing and editing of the draft. MP contributed to investigation, data curation, funding acquisition, resources, and supervision. SMB contributed to conceptualization, formal analysis, funding acquisition, methodology, supervision, reviewing and editing of the draft. All authors gave final approval of the version to be submitted.

## Conflicts of Interest

The authors declare no conflict of interest.

Correspondence and requests for materials should be addressed to SMB.

