## Supplementary table 1 for "A formula for calculating 30-item Geriatric Depression Scale (GDS-30) scores from the 15-item version (GDS-15)"

**Supplementary Table 1.** The interaction terms of age, sex and MMSE score with GDS-15 score in the estimation of GDS-30 score.

| **Variables** | **95% confidential interval** | **p-value** |
| --- | --- | --- |
| Age*GDS15 | [-0.01,0.01] | 0.95 |
| Sex*GDS15 | [-0.02, 0.37] | 0.08 |
| MMSE*GDS15 | [-0.02,0.05] | 0.46 |
